# Volatiles from low R:FR-treated maize plants increase the emission of herbivore-induced plant volatiles in their neighbors

**DOI:** 10.1101/2024.10.05.616768

**Authors:** Rocío Escobar-Bravo, Bernardus C.J. Schimmel, Matthias Erb

## Abstract

Low Red (R) to Far Red (FR) light ratios, a light signal associated with vegetation shade, can prompt intact maize (*Zea mays*) plants to constitutively emit more volatiles when exposed to herbivory-induced plant volatiles (HIPVs). Here we investigated how simulated shading affects priming responses in the context of volatile-mediated plant-plant interactions. Receiver maize plants were exposed to either constitutive volatile organic compounds (cVOCs) or HIPVs from emitter maize plants, while we manipulated R:FR light conditions of receivers or emitters. Priming responses in the receivers were then assessed by measuring real-time volatile emissions following simulated herbivory. We show that low R:FR light enhances HIPVs emissions in plants previously exposed to HIPVs from neighbours independently of the light conditions of emitters. We also demonstrate that both cVOCs and HIPVs emitted by maize grown under low R:FR amplify HIPVs emissions in their neighbours. This amplified response could not be explained by FR-mediated changes in the release of green leaf volatiles or terpenoids by emitters, thus suggesting the involvement of other VOCs. We conclude that volatile-mediated plant-plant interactions can be expected to become more intense in denser canopies due to light-mediated amplification of volatile emission and responsiveness.

## INTRODUCTION

Plants can sense the presence of neighbors through light, chemical, and mechanical cues (Huber et al., 2020; Ninkovic et al., 2021; Pantazopoulou et al., 2023). This ability to detect nearby plants helps them to assess their competitive environment, including the availability of light and soil nutrients, as well as the health and potential threats posed by pests and pathogens. By integrating these signals, plants can strategically allocate their resources towards growth or defense, thus enhancing their chances of survival.

Neighbor detection through light cues occurs via the photoreceptor-mediated perception of light spectral changes in the canopy. Blue (λ =400-500 nm) and red (R) (λ = 600-700 nm) light wavelengths are preferentially absorbed by photosynthetic pigments of plant leaves while far red (FR) light is mainly reflected or transmitted. In dense canopies, this results in a reduction in R:FR ratios, a light cue that can promote upward bending of the leaves (hyponasty) and stem elongation (Pierik and Ballaré 2021). These photomorphogenic changes help the plant to reach more illuminated areas in the canopy and outcompete their neighbors (Pierik and Ballaré 2021) In addition, FR photons can support growth by promoting photosynthesis in dicotyledonous and C3 and C4 monocotyledonous species, including maize (*Zea mays*), when added to a background of white light (Zhen and Bugbee 2020; Escobar□Bravo et al. 2024; Huber et al. 2024).

Beyond light, plants also perceive their environment through volatile organic compounds (VOCs) (Wang and Erb 2022). These compounds include volatile hormones (e.g., ethylene, methyl salicylate, and methyl jasmonate), terpenoids, benzenoids (e.g., indole), phenylpropanoids, and fatty acid-derived molecules such as green leaf volatiles (GLVs). Plant VOCs can be constitutively emitted and modulated by the plant interactions with the environment (Piesik et al. 2015; Escobar-Bravo et al. 2023). Changes in their emission therefore reflect the plant status and the abiotic and/or biotic stresses it encounters. Upon herbivory, for instance, plants can emit herbivory-induced plant volatiles (HIPVs) that repel the attacker, attract natural predators of the herbivore, and/or alter the defense status in non-attacked neighbors (Turlings and Erb 2018; Hu et al. 2021).

Some HIPVs, such as the green leaf volatiles (*Z*)-3-hexenal [(*Z*)-3-HAL], (*Z*)-3-hexen-1-ol [(*Z*)-3-HOL], or (*Z*)-3-hexenyl acetate [(*Z*)-3-HAC], can directly induce the emission of volatiles in intact plants and prime their volatile emissions (Farag and Pare 2002; Engelberth et al. 2004; Hu et al. 2019; Escobar□Bravo et al. 2024). Similarly, other HIPVs like indole, the monoterpene β-ocimene, and the homoterpene (*E*)-4,8-dimethyl-1,3,7-nonatriene (DMNT) can induce or prime plant defenses (Erb et al. 2015; Jing et al. 2021; Onosato et al. 2022). Defense priming can confer ecological benefits to plants, as the exposure to the priming stimulus prepares them to display a faster, stronger, and more lasting defense response when the actual herbivore or pathogen attack takes place (Conrath et al. 2015; Martinez-Medina et al. 2016). The mechanisms of volatile-mediated defense priming downstream of volatile perception have been amply characterized. Once HIPVs get into plant tissues, they induce the expression of biotic and abiotic stress-responsive genes in a Ca^2+^ dependent manner, including the activation of the jasmonic acid (JA) defense signaling pathway that controls the emission of HIPVs (Zebelo et al. 2012; Christensen et al. 2013; Engelberth et al. 2004; Hu et al. 2019; Ye et al. 2019; Jing et al. 2021; Aratani et al. 2023; Aguirre et al. 2023; Wang et al. 2023).

How VOC-mediated transfer of information between plants occurs in dense canopies is not well known. The FR-enriched light environment within vegetation canopies might play a crucial role in these interactions by altering (1) the emitter’s volatile emissions, (2) the ability of the receiver to perceive and respond to the volatile cue/s, or (3) both. For instance, Kegge et al. (2015) showed that barley (*Hordeum vulgare*) emits different volatiles under low R:FR ratios, influencing the carbon allocation in neighboring plants. Aguirre et al. (2023) reported that maize exposure to (*Z*)-3-HAC in the dark does not prime plant defenses, suggesting that light-mediated induction of stomatal opening is required for signal integration of (*Z*)-3-HAC in receiver plants. More recently, we have demonstrated that low R:FR light conditions can change the perception of HIPVs in maize plants (Escobar□Bravo et al. 2024). Our study showed that FR-light enrichment prompts intact maize plants to emit more volatiles when exposed to HIPVs from *Spodoptera littoralis*-infested neighbors or to (*Z*)-3-HAC alone. We also showed that low R:FR-treated plants pre-exposed to (*Z*)-3-HAC displayed stronger HIPVs emissions after simulated herbivory and, therefore, a stronger priming response. Yet, whether low R:FR-induced priming response also occurs in plants exposed to the whole bouquet of HIPVs from herbivore-infested neighbors was not determined. Furthermore, whether FR light enrichment can alter volatile-mediated interactions between maize plants by inducing changes in the VOCs emissions of emitter plants is unknown.

Here, we have investigated whether low R:FR affects HIPVs-induced priming responses in maize by altering (1) the volatile perception of the receivers, and/or (2) the volatile emissions of the emitters. For this, we first exposed plants to low R:FR ratios as a light cue for vegetation shade or high R:FR ratios as control light conditions, and we determined volatile emissions in simulated herbivory-induced plants that were previously exposed to constitutive volatile organic compounds (cVOCs) or HIPVs emissions from intact or *S. littoralis*-infested plants, respectively. Next, we conducted a multifactorial experiment where we tested whether HIPVs emitted by intact or simulated herbivory-induced maize plants growing under low or high R:FR conditions trigger different defense priming responses in the receivers, and whether these differences could be explained by changes in the GLVs emissions of the emitters. Our findings show that maize seedlings exposed to enriched FR conditions not only display a higher emission of HIPVs after exposure to volatile cues from neighbors, but they also emit volatiles that increase the emission of HIPVs in neighboring plants.

## METHODS & MATERIALS

### Plants

Experiments were conducted with the maize (*Zea mays*) inbred line B73. Seeds were sown in transparent cylindric plastic pots (11×4 cm) filled with commercial soil (Selmaterra, BiglerSamen, Switzerland) and wrapped with aluminium foil to prevent root exposure to light. Plants were grown in a ventilated greenhouse supplemented with artificial lighting under 50-70% relative humidity, 14/10 h light/dark photoperiod, and 14-22/10-14°C day/night temperatures.

### Insects

Eggs of *Spodoptera littoralis* were provided by Oliver Kindler (Syngenta, Stein, CHE) and the larvae were reared on artificial diet as described in (Maag et al. 2014). *S. littoralis* oral secretions were collected from third-to fourth-instar larvae as described in Hu *et al*., 2019 and stored at − 80°C until use.

### Light Treatments

Light treatments were conducted in a custom-made high-throughput phenotyping platform designed for volatile analyses previously described in Escobar□Bravo et al. (2024). Eleven-day old maize seedlings were exposed to low (∼0.5) or high (∼2.65) R:FR ratios (R, λ = 600-700 nm; FR, λ = 700-800 nm) by providing supplemental FR to the same white light background (120 ± 15 μmol m^−2^ s^−1^) (Supplemental Fig. S1) with a photoperiod of 16/8 h light/dark. This low R:FR corresponds to the ratios observed in high density maize canopies (Maddonni et al. 2002), where low PAR levels as the ones used in our experimental set-up are commonly observed (Xue et al. 2016). Light treatments were separated by white opaque curtains and started between 9:00 and 10:00 AM two days before the plant-plant interaction experiments.

### Plant-Plant Interaction Experiments

In the first experiment (Supplemental Fig. S2), we exposed receiver plants to low or high R:FR light conditions and the emitters to low R:FR light conditions. After two days, low R:FR-treated emitter plants were infested with ten second-instar *S. littoralis* larvae or left uninfested. Right after infestation, low and high R:FR-treated receiver plants were exposed to volatiles emitted by *S. littoralis-*infested or non-infested low R:FR treated plants. The chambers of emitter and receiver plants were connected using Teflon tubing. Clean air flow moved from the emitter’s chamber towards the receiver’s chamber. After 20 h, the Teflon tubing connecting emitters and receivers’ chambers were removed, and receiver plants were induced with simulated herbivory. Receiver’s volatile emissions were then recorded in time-series analysis using a PTR-TOF-MS system. In the second experiment (Supplemental Fig. S3), we employed the same set-up, but emitters were exposed to high R:FR light conditions.

In the third experiment (Supplemental Fig. S4), we conducted a multifactorial experiment where we exposed both emitter and receiver plants to high or low R:FR conditions for 2 days. Next, emitter and receivers’ chambers were connected via Teflon tubing as in previous experiments and the emitter plants were immediately induced with simulated herbivory or left intact. By using simulated herbivory, we standardized the leaf damage and minimized differences in herbivore-associated damage patterns between low and high R:FR-treated emitter plants. After 6 h, the Teflon tubing connecting emitters and receivers’ chambers was removed, and all the receiver plants were induced with simulated herbivory. Although the duration of the exposure to volatiles was shorter than in Experiments 1 and 2, our previous work has shown that HIPVs emissions in simulated herbivory induced maize plants steadily increases and reaches their maximum over the first six hours after induction, and that exposure to GLVs for 6 h can prime volatile emissions (Escobar□Bravo et al. 2024). Receiver’s volatile emissions were recorded in time-series analysis using a PTR-TOF-MS system. This experiment was repeated three times using three replicates per combined treatment each time. Data from the three repetitions were pooled prior to the statistical analysis.

### PTR-TOF-MS Volatile Emission Analysis

Plant volatile emission was determined using the automated high-throughput real-time phenotyping platform and the methods described in Escobar□Bravo et al. (2024)

### Simulated Herbivory

Two leaves (leaf 2 and 3 from the bottom) per plant were wounded over an area of ca. 0.5 cm^2^ parallel to the central vein using a haemostat followed by the application of 10 μL of *S. littoralis* larval oral secretions (diluted 1:1 in autoclaved distilled water) in each wounding site. This treatment induces plant defence responses comparable to real herbivory (Erb et al. 2009).

### Statistical Analysis

Data analysis was conducted in R version 4.0.4 (R core Team, 2016). Effects of (1) sampling time, (2) light receiver, (3) light emitter, (4) volatiles from the emitter, and (5) their interactions on volatile emissions in the time course experiments (i.e., Figures 1, 2, 3, and 5) were tested in linear mixed-effects models (LME) using the nlme package. Individual plants were included in the model as a random factor and a correlation structure when autocorrelation among residuals was found to be significant (*p* < 0.05). Fitted models were subjected to type III analyses of variance (ANOVAs) to produce a summary of the F- and p statistics (car package) (Fox et al. 2012). When needed, data from volatiles signatures were log10 or squared root transformed prior analysis to correct for heteroscedasticity. Cumulative emission of the sum of all volatile’s signatures depicted in Fig. 4 was analysed by using a mixed-effects model where we the replication of the experiment was included as random factor. Cumulative emission of individual volatiles over time depicted in Fig. S5 and S6 was analysed by using linear models. Differences among treatments were tested by the calculation of estimated marginal means (EMMs) followed by pairwise comparisons using Tukey’s Honest Significant Difference (HSD) test (emmeans package; Lenth, 2021).

**Figure 1.**
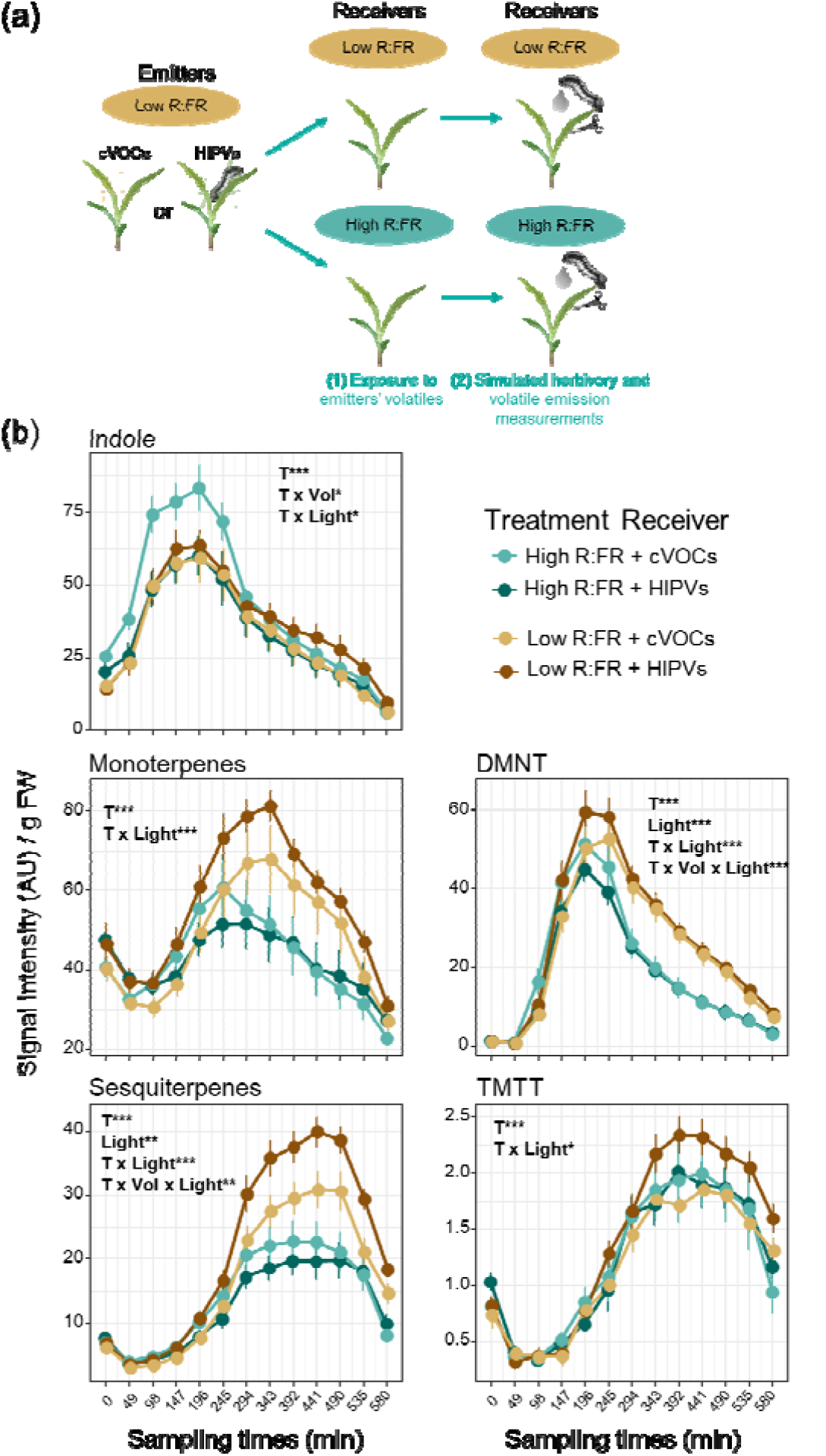
Low R:FR enhances priming of monoterpenes, sesquiterpenes and TMTT emissions in maize plants pre-exposed to HIPVs from low R:FR treated plants. (a) Schematic overview of the emitters and receivers’ treatments. (b) Time-series analysis of volatile emissions (mean ± SEM, *n* = 6) collected from receiver plants pre-exposed to cVOCs or HIPVs from low R:FR-treated plants during 20 h and followed by simulated herbivory (wounding + oral secretions from *S. littoralis*). The effects of sampling time (T), light treatment of receivers (Light), volatiles from emitters (Vol), and their interactions on receivers’ volatile emissions were tested using linear mixed-effects models. Statistically significant effects are shown in each graph (\**p* < 0.05, ** *p* < 0.001, *** *p* < 0.001). DMNT and TMTT stand for the homoterpenes (*E*)-4,8-dimethyl-1,3,7-nonatriene and (*E, E*)-4,8,12-trimethyltrideca-1,3,7,11-tetraene, respectively. AU refers to arbitrary units.

**Figure 2.**
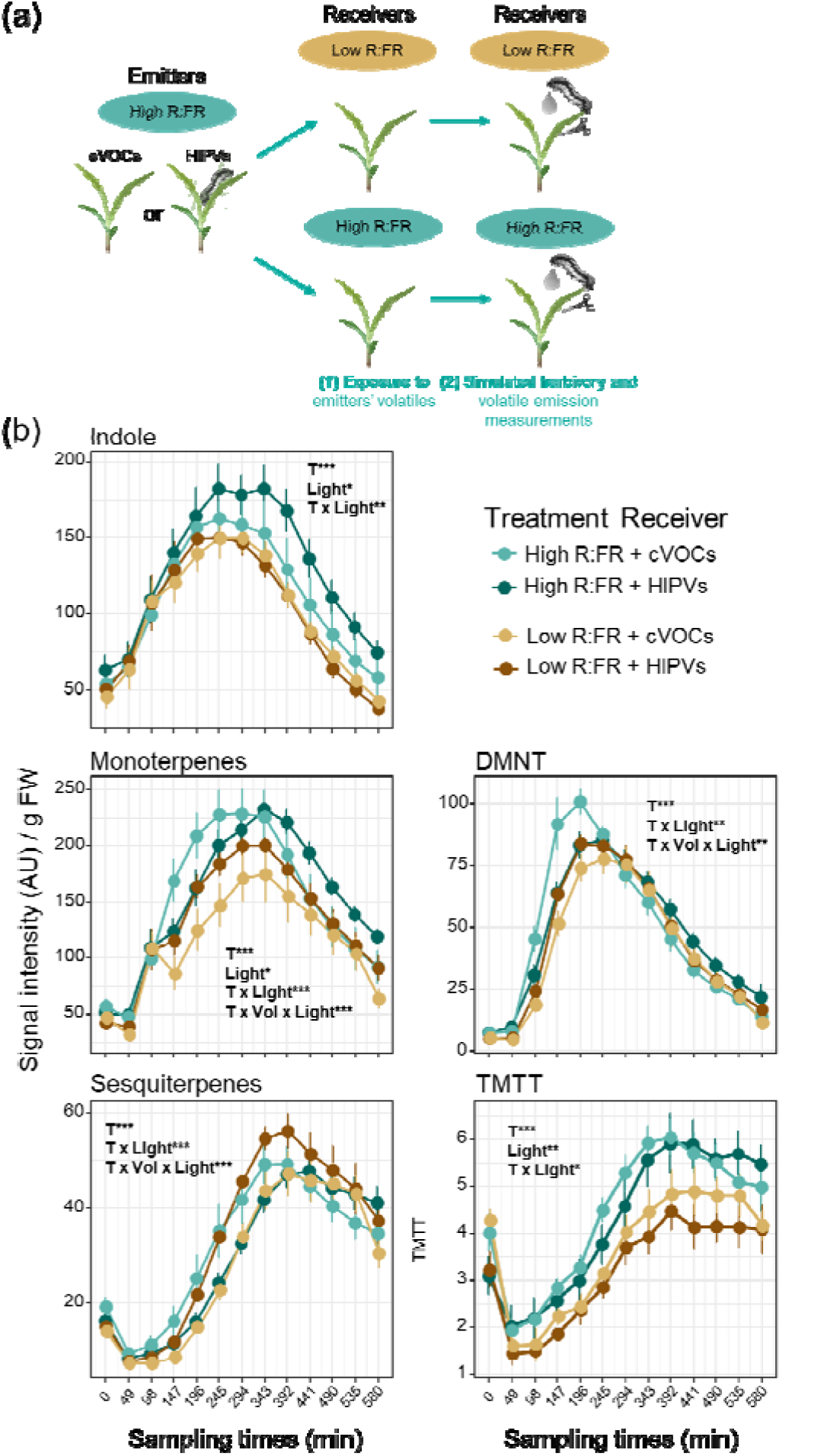
Low R:FR enhances priming of monoterpenes and sesquiterpenes emissions in maize plants pre-exposed to HIPVs from high R:FR treated plants. (a) Schematic overview of the emitters and receivers’ treatments. (b) Time-series analysis of volatile emissions (mean ± SEM, *n* = 6) collected from receiver plants pre-exposed to cVOCs or HIPVs from low R:FR-treated plants during 20 h and followed by simulated herbivory (wounding + oral secretions from *S. littoralis*). The effects of sampling time (T), light treatment of receivers (Light), volatiles from emitters (Vol), and their interactions on receivers’ volatile emissions were tested using linear mixed-effects models. Statistically significant effects are shown in each graph (\**p* < 0.05, ** *p* < 0.001, *** *p* < 0.001). DMNT and TMTT stand for the homoterpenes (*E*)-4,8-dimethyl-1,3,7-nonatriene and (*E, E*)-4,8,12-trimethyltrideca-1,3,7,11-tetraene, respectively. AU refers to arbitrary units.

**Figure 3.**
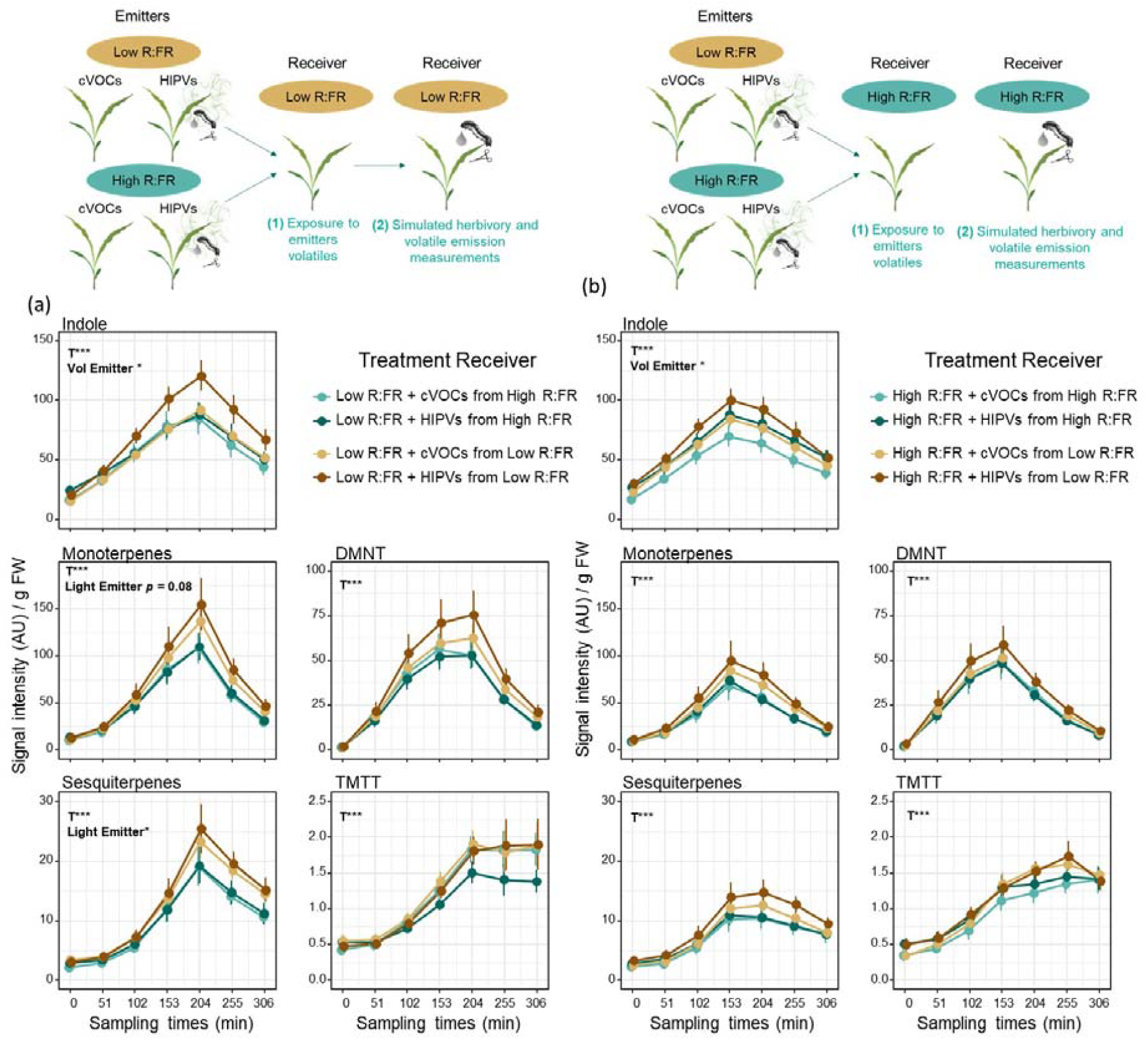
Volatiles emitted by low R:FR-treated plants enhance emissions of HIPVs in low R:FR-treated plants. Time-series analysis of volatile emissions (mean ± SEM, *n* = 9) collected from **(a)** low R:FR-treated or **(b)** high R:FR-treated receiver plants pre-exposed to cVOCs or HIPVs from low or high R:FR-treated plants during 6 h and followed by simulated herbivory (wounding + oral secretions from *S. littoralis*). Pooled data from three independent experiments are shown. The effects of sampling time (T), light treatment of emitters (Light Emitter), volatiles from emitters (Vol Emitter), and their interactions on receivers’ volatile emissions were tested using linear mixed-effects models. Statistically significant effects are shown in each graph (\**p* < 0.05, ** *p* < 0.001, *** *p* < 0.001). DMNT and TMTT stand for the homoterpenes (*E*)-4,8-dimethyl-1,3,7-nonatriene and (*E, E*)-4,8,12-trimethyltrideca-1,3,7,11-tetraene, respectively. AU refers to arbitrary units.

**Figure 4.**
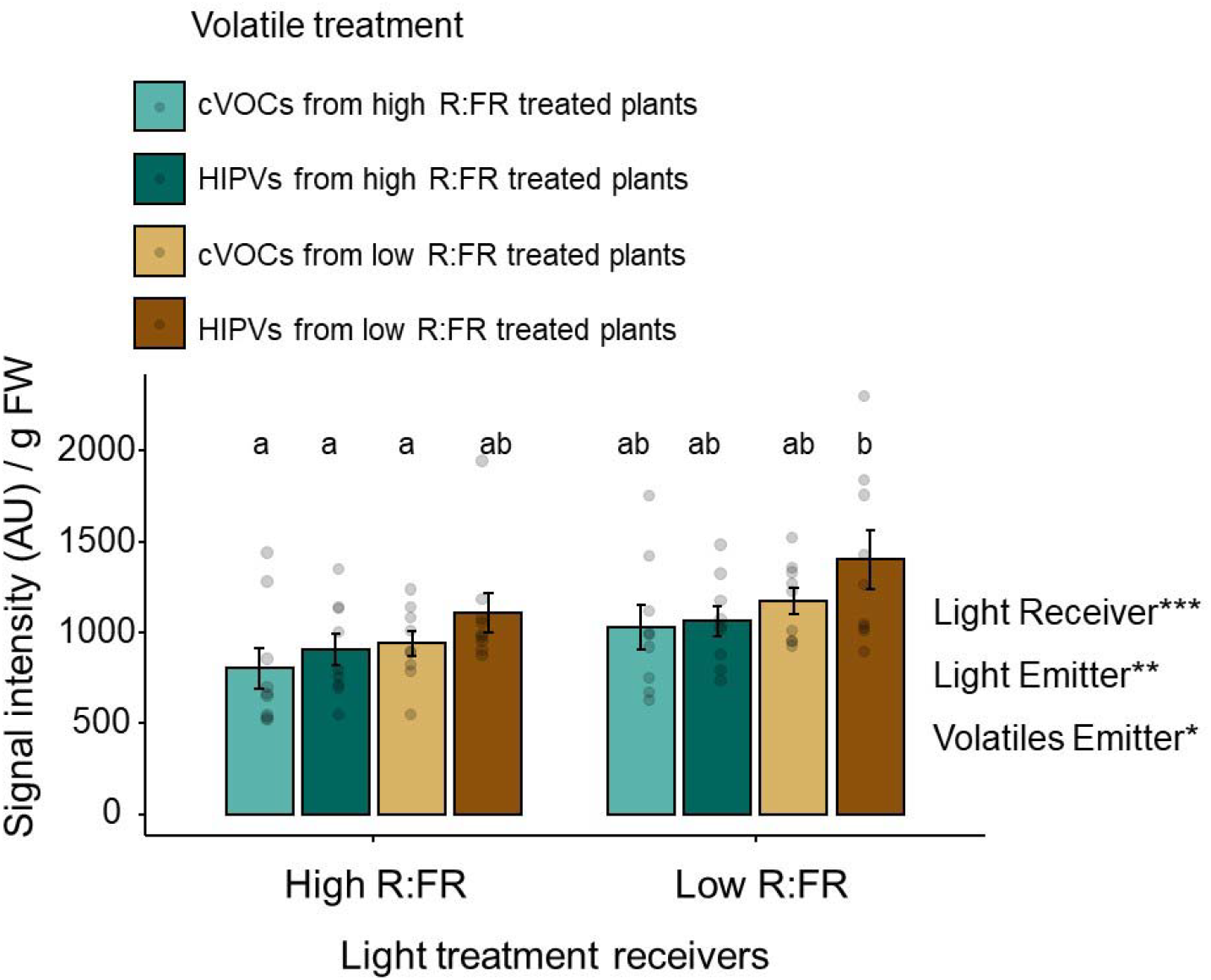
Sum (mean ± SEM, *n* = 9) of the emission of all the volatiles signatures (indole, monoterpenes, sesquiterpenes, DMNT, and TMTT) measured over a period of 5 h upon simulated herbivory induction in receiver plants. Low or high R:FR-treated receiver plants were pre-exposed to cVOCs or HIPVs from low or high R:FR-treated plants during 6 h followed by simulated herbivory (wounding + oral secretions from *S. littoralis*).The effects of light treatment of the receiver (Light Receiver), light treatment of the emitter (Light Emitter), the volatiles from the emitters (Volatiles Emitter), and their interactions on receivers’ volatile emissions were tested using mixed-effects models followed by the calculation of estimated marginal means (EMMs) and pairwise comparisons using Tukey’s Honest Significant Difference (HSD) test. Statistically significant effects are shown in the graph (\**p* < 0.05, \*\**p* < 0.01, \*\*\**p* < 0.001). Different letters denote significant differences among groups at *p* < 0.05.

**Figure 5.**
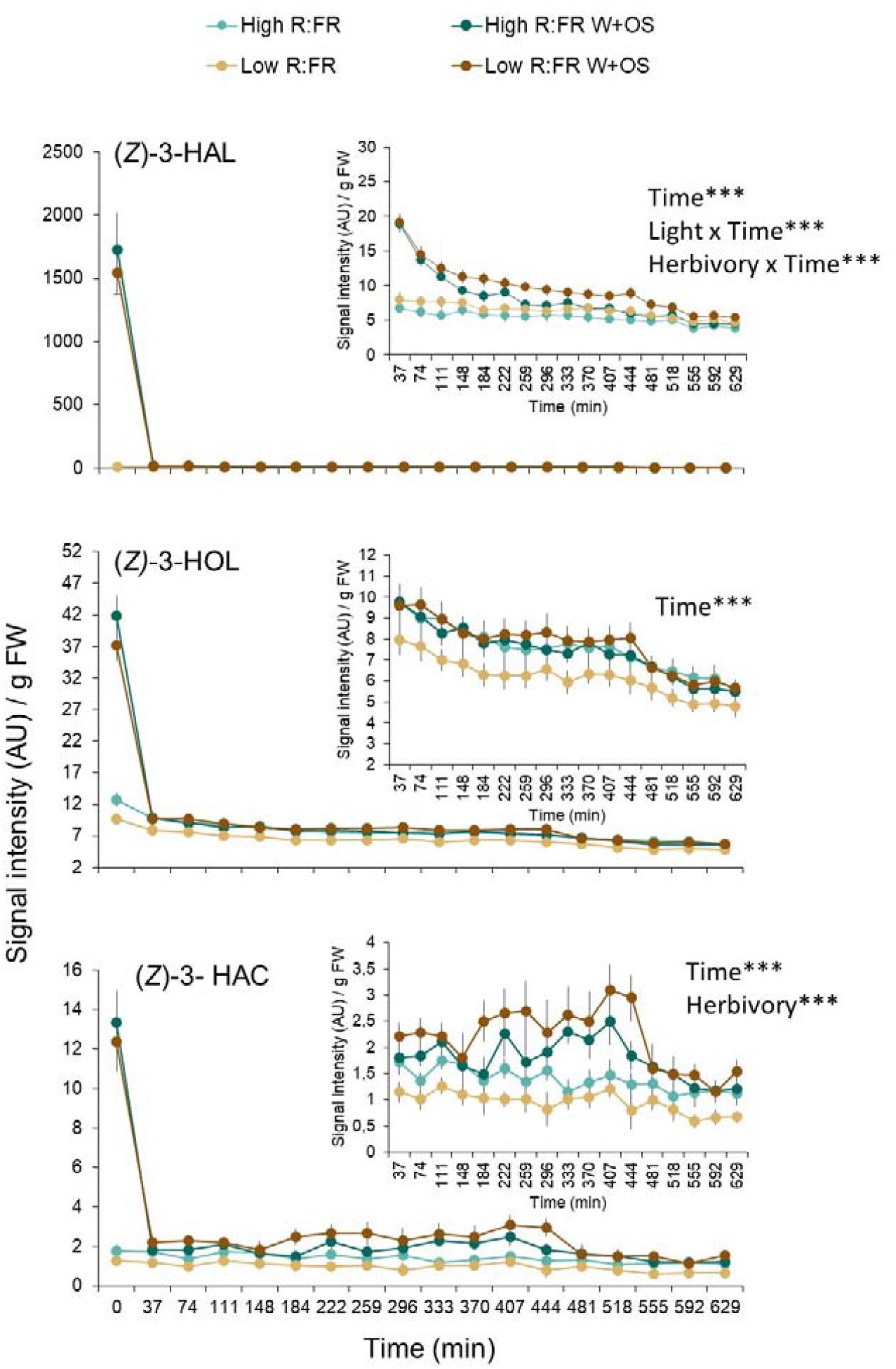
Time series analysis of green leaf volatile emissions (mean ± SEM, *n* = 8) determined by PTR-TOF-MS at intervals of 37 min in high and low R:FR-treated B73 maize (*Zea mays*) plants after simulated herbivory. Volatile measurements started at 12:30 pm, 10 min after simulated herbivory treatment (wounding and application of *Spodoptera littoralis* oral secretions; W+OS). The effects of sampling time (Time), light treatment (Light), simulated herbivory (Herbivory) and their interactions on volatile emission were tested using linear mixed-effects models. Plant unit was included as the random intercept as well as a correlation structure when autocorrelation among residuals was found significant (*p* < 0.05). Statistically significant effects are indicated in each graph. *** *p* < 0.001.

## RESULTS

### Low R:FR Light Enhances HIPVs-Mediated Priming of Volatile Emissions in Maize

Low R:FR light ratios can increase the emission of volatiles in intact maize plants when these are exposed to HIPVs emitted by low R:FR-treated neighbors (Escobar□Bravo et al. 2024). To determine whether low R:FR and pre-exposure to HIPVs emitted by herbivore-infested neighbors affects subsequent maize responses to herbivory, we conducted a series of plant-plant interactions experiments.

In the first experiment, we exposed 11-day old B73 maize seedlings to low or high R:FR light conditions for two days and measured their volatile emissions after exposure to volatiles emitted by *S. littoralis-*infested or non-infested low R:FR treated plants followed by induction with wounding and the application of *S. littoralis* oral secretions (Fig. S2 and Fig. 1ab). Upon simulated herbivory, emission of all the volatile signatures increased over time (Fig. 1b). This induction was stronger under low R:FR irrespective of the pre-exposure to emitters’ volatiles. Emissions of monoterpenes, sesquiterpenes, DMNT, and TMTT [(*E, E*)-4,8,12-trimethyltrideca-1,3,7,11-tetraene] were higher in low R:FR treated plants compared to high R:FR treated plants, except for indole. Light also influenced maize responses to volatiles. The emissions of sesquiterpenes, monoterpenes, TMTT, and DMNT were higher in low R:FR-treated plants exposed to HIPVs from herbivore-infested plants, whereas no induction, or even the opposite pattern, was observed in high R:FR-treated plants. Cumulative emissions of monoterpenes and sesquiterpenes were higher in low R:FR treated plants pre-exposed to HIPVs in comparison to their controls pre-exposed to cVOCs (Fig. S5)

In the second experiment, we used the same set-up, but plants were instead exposed to volatiles emitted by high R:FR treated plants that were either infested with *S. littoralis* larvae or non-infested (Fig S3, and Fig. 2). Emission of all the volatiles increased over time upon simulated herbivory in low and high R:FR treated plants. Unlike the first experiment, plants exposed to low R:FR and volatiles from high R:FR treated plants emitted less indole, monoterpenes, DMNT and TMTT than plants exposed to high R:FR and the same volatiles (Fig. 2 and Fig. S6). Prior exposure to volatiles from high R:FR treated plants infested with *S. littoralis* larvae slightly enhanced the emission of monoterpenes and sesquiterpenes in low R:FR-treated plants upon simulated herbivory. In plants irradiated with high R:FR ratios, prior exposure to HIPVs slightly decreased the emission of monoterpenes, sesquiterpenes, and DMNT over time, and the opposite was observed for indole. Similar patterns were found when the cumulative emissions of individual volatiles were analyzed (Fig. S6). Low R:FR treated plants emitted less indole, monoterpenes and TMTT than high R:FR-treated plants upon exposure to emitter’s volatiles and simulated herbivory.

These results suggest that low R:FR can enhance plant responses to HIPVs, but this induction seems to be stronger when plants were exposed to volatiles emitted by neighbors growing under low R:FR ratios.

### Volatiles Emitted by Low R:FR Treated Maize Plants Increase the Emission Of Herbivore-Induced Plant Volatiles in Receiver Plants

To further test whether maize plants respond differently to volatiles emitted by plants under low R:FR ratios (i.e., simulated shade) and control for potential HIPVs differences arising from herbivore feeding patterns, we conducted a factorial experiment where high and low R:FR-treated plants were exposed to volatiles emitted by low or high R:FR treated plants that were either induced with simulated herbivory or left intact (Supplemental Fig. S4). This set-up allowed us to compare in the same experiment the magnitude of the priming induction when the emitters differ in their light environment conditions but similar herbivory induction. After 6 h of exposure to constitutive or herbivory-induced volatiles from emitters, receiver plants were induced with simulated herbivory and their volatile emissions were determined by PTR-TOF-MS in time series analyses.

For the sake of clarity, we present the results of this experiment in two panels grouped by the light treatment of the receiver (Fig. 3). Maize plants exposed to low R:FR responded differently to volatiles emitted by low and high R:FR-treated plants (Fig. 3a). Upon simulated herbivory, monoterpenes, sesquiterpenes and linalool emissions were higher when low R:FR-treated plants were exposed to volatiles emitted by low R:FR-treated plants than by emitter plants irradiated with high R:FR (Fig. 3a). This induction was statistically significant for sesquiterpenes, and it was independent of the herbivory treatment of the emitters. In addition, indole emissions were increased in low R:FR-treated plants when they were pre-exposed to volatiles from not-wounded low R:FR-treated plants.

Receiver plants exposed to high R:FR ratios and volatiles emitted by low R:FR-treated plants showed a slight increase in the emission of monoterpenes and sesquiterpenes, but this was not statistically significant for any of the volatile signatures (Fig. 3b).

Further analysis on the sum of all the volatiles emitted over the sampling period in both data sets showed that (1) low R:FR-treated plants overall emitted more volatiles after exposure to volatiles from emitters and subsequent simulated herbivory and (2) volatiles emitted by intact (cVOCs) and herbivory-induced (HIPVs) low R:FR-treated plants enhance the emission of HIPVs in neighboring plants (Fig. 4).

### Low R:FR Slightly Increases (Z)-3-HAL Emission but not (Z)-3-HOL, and (Z)-3-HAC

Green leaf volatiles such as (*Z*)-3-HAL, (*Z*)-3-HOL, and (*Z*)-3-HAC, and indole are HIPVs emitted by maize that can primed defenses in neighboring plants (Farag and Pare 2002; Engelberth et al. 2004; Erb et al. 2015). Yet, our previous study has shown that indole emissions in low R:FR treated plants do not differ from high R:FR-treated plants upon simulated herbivory (Escobar□Bravo et al. 2024). To further determine whether FR light affect the emission of GLVs, we compared (*Z*)-3-HAL, (*Z*)-3-HOL, and (*Z*)-3-HAC emissions in simulated herbivory-induced and unwounded plants that were previously exposed to low or high R:FR conditions for 2 days (Fig. 5).

Simulated herbivory significantly induced the emissions of (*Z*)-3-HAL, (*Z*)-3-HOL, and (*Z*)-3-HAC in both low and high R:FR treated plants (Fig. 5). Low R:FR did not significantly affect (*Z*)-3-HOL, and (*Z*)-3-HAC emissions in unwounded and simulated herbivory-induced plants (Fig. 5bc). (*Z*)-3-HAL emission levels were higher in low R:FR-treated plants compared to high R:FR treated plants (Fig. 5a).

## DISCUSSION

Our study shows that maize plants growing under FR-enriched light conditions (low R:FR ratios) not only display a higher emission of HIPVs after exposure to volatile cues from neighbors, but they also emit volatiles that increase the emission of HIPVs in neighboring plants. These findings contribute to unraveling the complexity of volatile-mediated interplay between plants in vegetation canopies where shade-associated light cues might play a key role.

Our previous study demonstrated that low R:FR enhances HIPVs emissions in plants exposed to (*Z*)-3-HAC alone followed by simulated herbivory (Escobar-Bravo et al., 2024). Here we show that maize plants exposed to low R:FR also display a stronger priming response upon pre-exposure to the HIPVs emitted by herbivore-infested plants. We hypothesize that this response might be partially explained by FR-mediated positive effects on stomata opening and photosynthesis (Zhen and Bugbee 2020; Escobar□Bravo et al. 2024; Huber et al. 2024). As stomata are the entry ports of VOCs into intact plants (Aratani et al. 2023) and modulate indole and (*Z*)-3-HAL-mediated priming responses in maize (Aguirre et al. 2023), changes in stomatal aperture might have affected maize perception and responses to neighbor’s volatiles. In other words, FR-treated plants might take up more volatiles than high R:FR-treated plants, resulting in a stronger priming stimulus. Alternatively, FR positive effects on photosynthesis and stomatal opening might have increased the availability of precursors for volatile biosynthesis (Arimura et al. 2008) and/or the release rate of these volatiles. Further studies are needed to determine the mechanisms involved in FR-mediated modulation of plant perception and responses to HIPVs.

Next, we showed that maize plants emit more HIPVs after pre-exposure to constitutive or herbivory-induced volatiles from low R:FR-treated plants. A possible explanation for this phenomenon is that a higher emission of volatiles by low R:FR-treated plants might trigger a stronger priming response in the receiver. For instance, exposure to increasing concentrations of GLVs has been shown to parallel increases in plasma membrane potential depolarization and cytosolic calcium concentrations in plant leaves (Zebelo et al. 2012; Aratani et al. 2023). These cellular responses are associated with the activation of JA defense-related genes (Aratani et al. 2023) and might be part of the priming stimulus in the receiver plant. Our analysis of GLV emissions in intact plants and plants subjected to simulated herbivory showed slightly higher levels of hexenal in low R:FR-treated plants compared to the high R:FR treatment, but no differences in (*Z*)-3-HAC and (*Z*)-3-HOL emissions. Thus, it is unlikely that these differences are responsible for the increase in HIPVs observed in receiver plants.

Additionally, our previous study revealed that an enriched FR light environment enhances the emission of monoterpenes and the homoterpene DMMT in herbivory-induced B73 maize plants (Escobar□Bravo et al. 2024). Yet, in the same experiments, we did not observe significant differences in constitutive volatile emissions in unwounded FR-treated plants compared to high R:FR light conditions (Escobar□Bravo et al. 2024). As both unwounded and herbivory-induced low R:FR treated plants enhanced the volatile emissions in receiver plants, they might emit a common volatile signal that was not detected in our PTR-TOF-MS system. This would be the case of volatile compounds that cannot be ionized by the primary ion (H_3_O^+^) used in our system. A candidate signal could be the volatile hormone ethylene. Ethylene emission is induced by low R:FR light conditions in several plant species, including *Arabidopsis thaliana* and *Nicotiana tabacum*, playing an important role in the development of shade avoidance traits (Pierik et al. 2003, 2004; Kegge et al. 2013). Furthermore, ethylene exposure has been shown to increase the emission of HIPVs in maize plants when these were challenged with simulated herbivory (Schmelz et al. 2003) or (*Z*)-3-HOL exposure (Ruther and Kleier 2005). Further analysis determining ethylene emissions in low R:FR-treated maize plants together with the artificial manipulation of ethylene signaling could shed light on the role of this volatile hormone in the stronger volatile-mediated priming response under enriched FR light conditions.

Finally, our data showed that while 20 h of exposure to HIPVs from plants irradiated with low R:FR ratios and infested with *S. littoralis* larvae induced a clear priming response in low R:FR treated plants, shorter exposure (6 h) to HIPVs from simulated herbivory-induced plants did not. We hypothesize that exposure time and differences in GLVs emission patterns between real and simulated herbivory might explain these differences and this would need further investigation.

In summary, our findings suggest an enhanced transmission of volatile signals within maize canopies via FR-mediated effects on the emitter’s volatile emissions and the perception of these volatiles by the receiver. Additional experiments are needed to determine (1) whether this translates into a larger spatial distribution of warning volatile signals in maize fields, (2) if this information transfer is species-specific, and (3) what are the benefits and tradeoffs of such a phenomenon. Under field conditions, for instance, the propagation of volatile signals involving several plants located at different distances from the emitter might be affected by the diffusion properties and breakdown of volatile compounds after release (Niinemets et al. 2014; Schuman 2023), as well as by their absorption and reabsorption by the plant/s closer to the volatile source (Himanen et al. 2010; Sugimoto et al. 2014, 2023). In addition, maize plants might be exposed to other plant-competition cues such as limited water and nutrient resources, and root-derived exudates from neighboring plants (Pierik et al. 2013; Wang et al. 2021) that might affect volatile emissions and perception. For instance, drought can modulate herbivory-induced (Catola et al. 2018; Lin et al. 2022; Vázquez□González et al. 2022) and constitutive volatile emissions (Jin et al. 2021), the latter affecting the drought tolerance in neighboring plants (Jin et al. 2021). Finally, we hypothesize that an enhanced transmission of volatile priming signals in maize fields might increase HIPVs emission at the plant community level which can boost the attraction of the natural enemies of herbivores (Aartsma et al. 2017). This might be especially relevant in dense monocultures, where the limited air movement within a dense canopy can reduce the dispersion of plant volatile compounds, potentially leading to volatile entrapment. Thus, our study not only provides a better understanding of the environmental modulation of the transmission of volatile priming signals in an agronomic relevant species but also sets the basis to further determine their ecological functions.

## ACKNOWLEDGEMENTS

The work of BCJS was supported by the Marie Sklodowska-Curie Action Individual Fellowship (European Union Horizon 2020, Grant Nr. 794947). This work was supported by the Velux Foundation (Grant Nr. 1231), the Swiss National Science Foundation (Grant. Nr. 200355), the State Secretariat for Education, Research, and Innovation SERI (Project CANWAS) and the University of Bern.

## AUTHORS CONTRIBUTION

ME conceived the study. ME, RE-B, BCJS designed experiments. RE-B and BCJS conducted the experiments. RE-B and BCJS performed the PTR-TOF-MS measurements and analyses. RE-B analyzed the data. RE-B, BCJS, and ME interpreted the data. RE-B wrote the manuscript, and all the authors contributed to revisions.

## DATA AVAILABILITY

The data generated for this manuscript and the code to reproduce the statistical analysis can be downloaded from GitHub (https://github.com/RocioEscobarBravo/Escobar-Bravo-et-al.-JCE).

## SUPPLEMENTARY MATERIAL

**Figure S1**. Light spectral composition determined in far-red supplementation experiments.

**Figure S2**. Detailed schematic overview of the experimental set-up in Fig. 1

**Figure S3**. Detailed schematic overview of the experimental set-up in Fig. 2.

**Figure S4**. Detailed schematic overview of the experimental set-up in Fig. 3 and Fig. 4.

**Figure S5**. Cumulative emission of individual volatiles in Fig. 1.

**Figure S6**. Cumulative emission of individual volatiles in Fig. 2.

