## Supplementary material for "Volatiles from low R:FR-treated maize plants increase the emission of herbivore-induced plant volatiles in their neighbors"

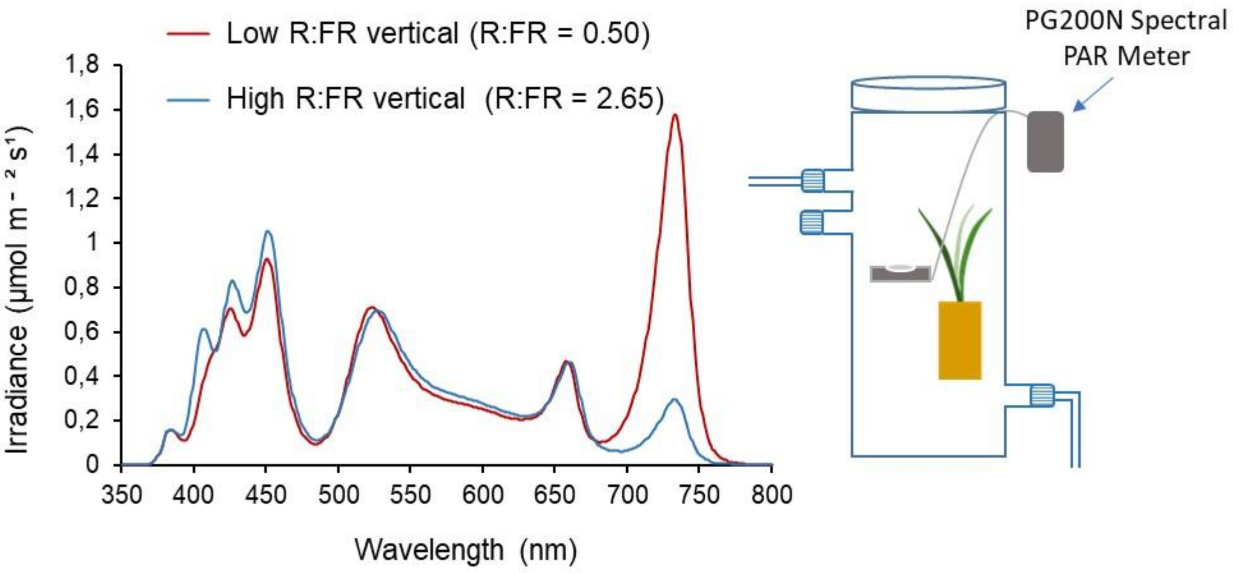


**Figure S1. Light spectral composition determined in far-red (FR) supplementation experiments using a PG200N Spectral PAR Meter (UPRtek).** Light spectra were vertically measured inside two representative glass chambers where 11-day-old maize (*Zea mays*) plants were exposed to low or high R:FR light conditions. Photosynthetically active radiation values under high or low R:FR light conditions in these two glass chambers were 121 and 111 µmol $m^{-2}$ $s^{1}$ respectively. Averaged PAR levels among experimental replicates for both low and high R:FR light treatments were the same, i.e., 120 ± 15 µmol $m^{-2}$ $s^{1}$. The legend indicates the ratio between Red (R: λ 600-700 nm) and Far Red (FR: λ 700-800 nm) light.

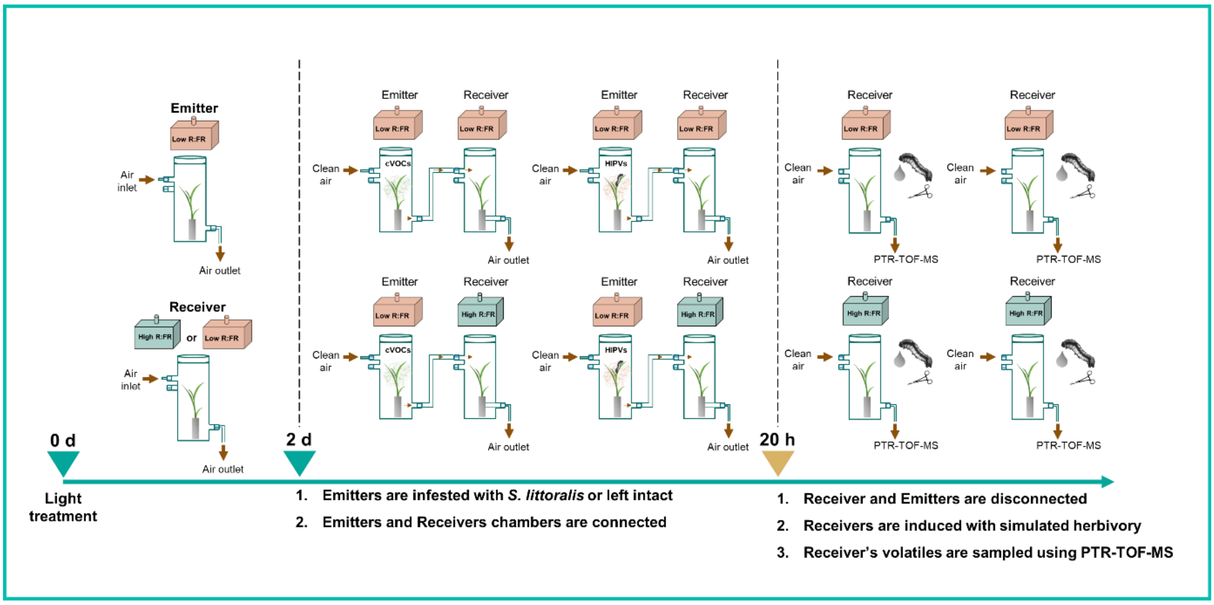


**Figure S2. Detailed schematic overview of the experimental set-up in Fig. 1.** Eleven-day old `B73´ maize (*Zea mays*) plants were exposed to low or high R:FR light conditions by modulating FR levels. After two days, receiver plants growing under low or high R:FR were exposed to constitutive volatiles (cVOCs) or *S. littoralis* herbivory-induced plant volatiles (HIPVs) from emitter plants exposed to low R:FR. Emitter and Receiver plants were connected via Teflon tubing. After 20 h, emitters and receivers' chambers were disconnected, and all the receiver plants were induced with simulated herbivory (wounding + oral secretions from *S. littoralis*). Volatile emissions in induced receivers were measured by PTR-TOF-MS.

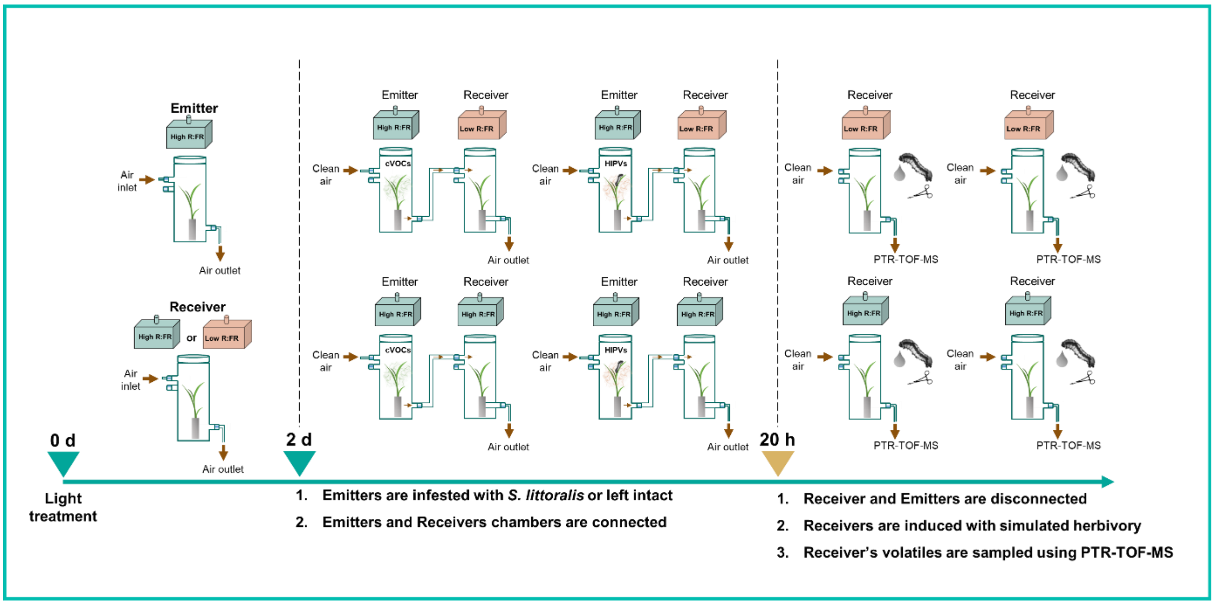


**Figure S3. Detailed schematic overview of the experimental set-up in Fig. 2.** Eleven-day old `B73´ maize (*Zea mays*) plants were exposed to low or high R:FR light conditions by modulating FR levels. After two days, receiver plants growing under low or high R:FR were exposed to constitutive volatiles (cVOCs) or *S. littoralis* herbivory-induced plant volatiles (HIPVs) from emitter plants exposed to low R:FR. Emitter and Receiver plants were connected via Teflon tubing. After 20 h, emitters and receivers' chambers were disconnected, and all the receiver plants were induced with simulated herbivory (wounding + oral secretions from *S. littoralis*). Volatile emissions in induced receivers were measured by PTR-TOF-MS.

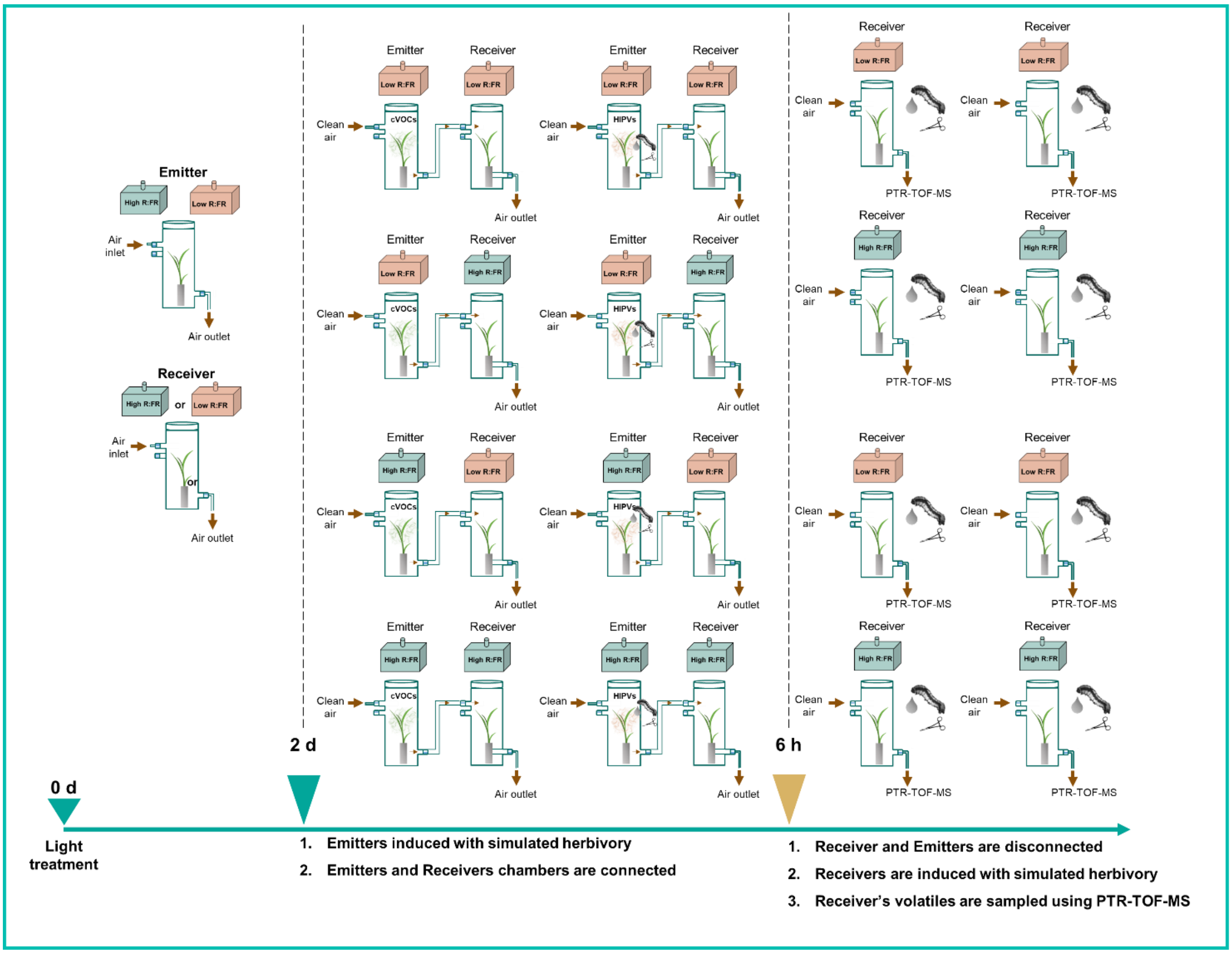


**Figure S4. Detailed schematic overview of the experimental set-up in Fig. 3 and Fig. 4.** Eleven-day old `B73´ maize (*Zea mays*) plants were exposed to low or high R:FR light conditions by modulating FR levels. After two days, receiver plants growing under low or high R:FR were exposed to constitutive volatiles (cVOCs) or herbivory-induced plant volatiles (HIPVs) from emitter plants exposed to low or high R:FR. Emitter and Receiver plants were connected via Teflon tubing. After 6 h, emitters and receivers' chambers were disconnected, and all the receiver plants were induced with simulated herbivory (wounding + oral secretions from *S. littoralis*). Volatile emissions in induced receivers were measured by PTR-TOF-MS.

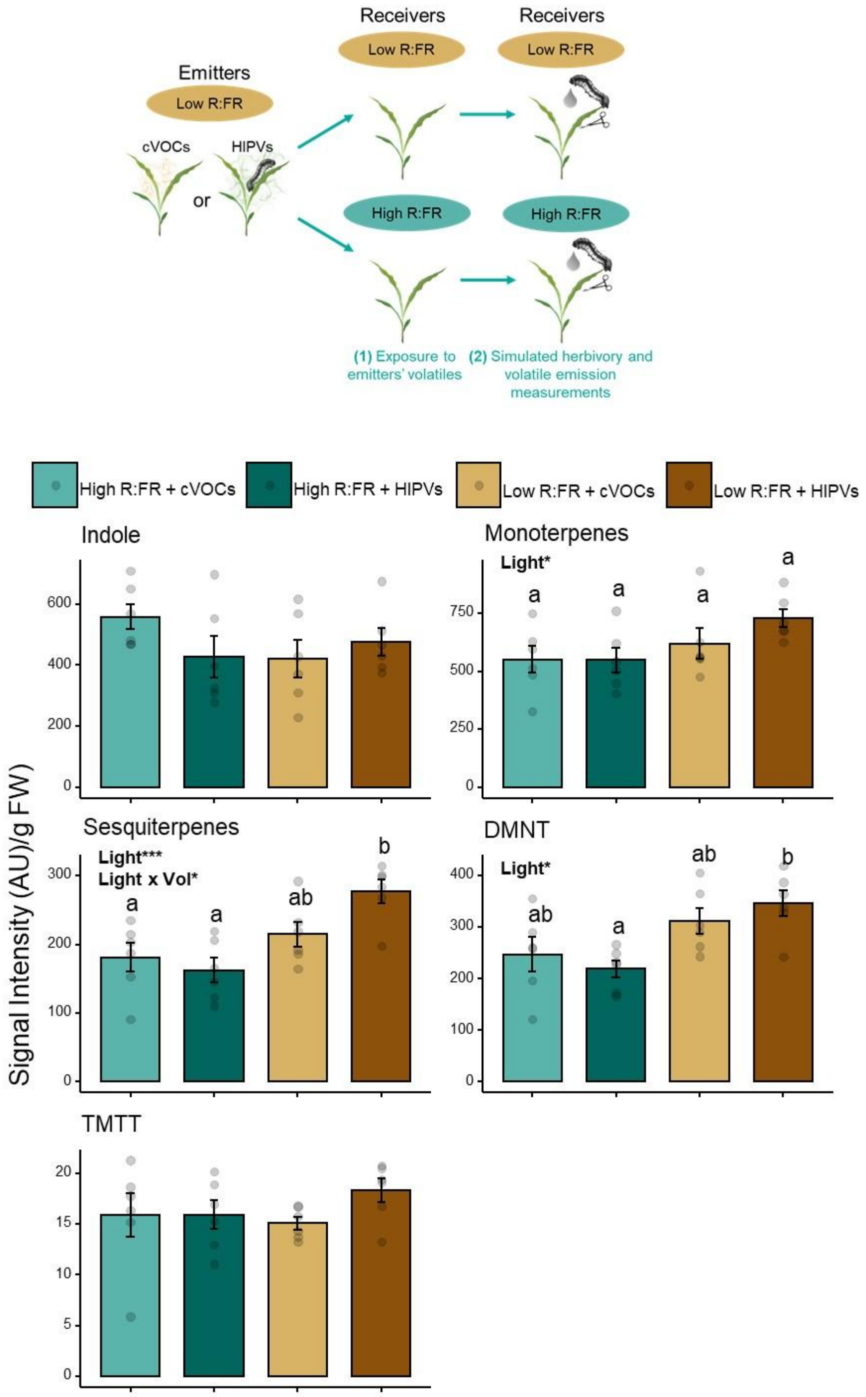


**Figure S5.** Sum (mean ± SEM, $n$ = 6) of the emission of individual volatiles signatures measured over a period of 12 h upon simulated herbivory in receiver plants. Low or high R:FR-treated receiver plants were pre-exposed to cVOCs or HIPVs from low R:FR-treated plants during 20 h followed by simulated herbivory (wounding + oral secretions from *S. littoralis*).The effects of light treatment of the receiver (Light), the volatiles from the emitters (Vol), and their interactions on receivers' volatile emissions were tested using linear models followed by the calculation of estimated marginal means (EMMs) and pairwise comparisons using Tukey's Honest Significant Difference (HSD) test. Statistically significant effects are shown in the graph (*$p < 0.05$, ***$p < 0.001$). Different letters denote significant differences among groups at $p < 0.05$. DMNT and TMTT stand for the homoterpenes (*E*)-4,8-dimethyl-1,3,7-nonatriene and (*E*, *E*)-4,8,12-trimethyltrideca-1,3,7,11-tetraene, respectively. AU refers to arbitrary units.

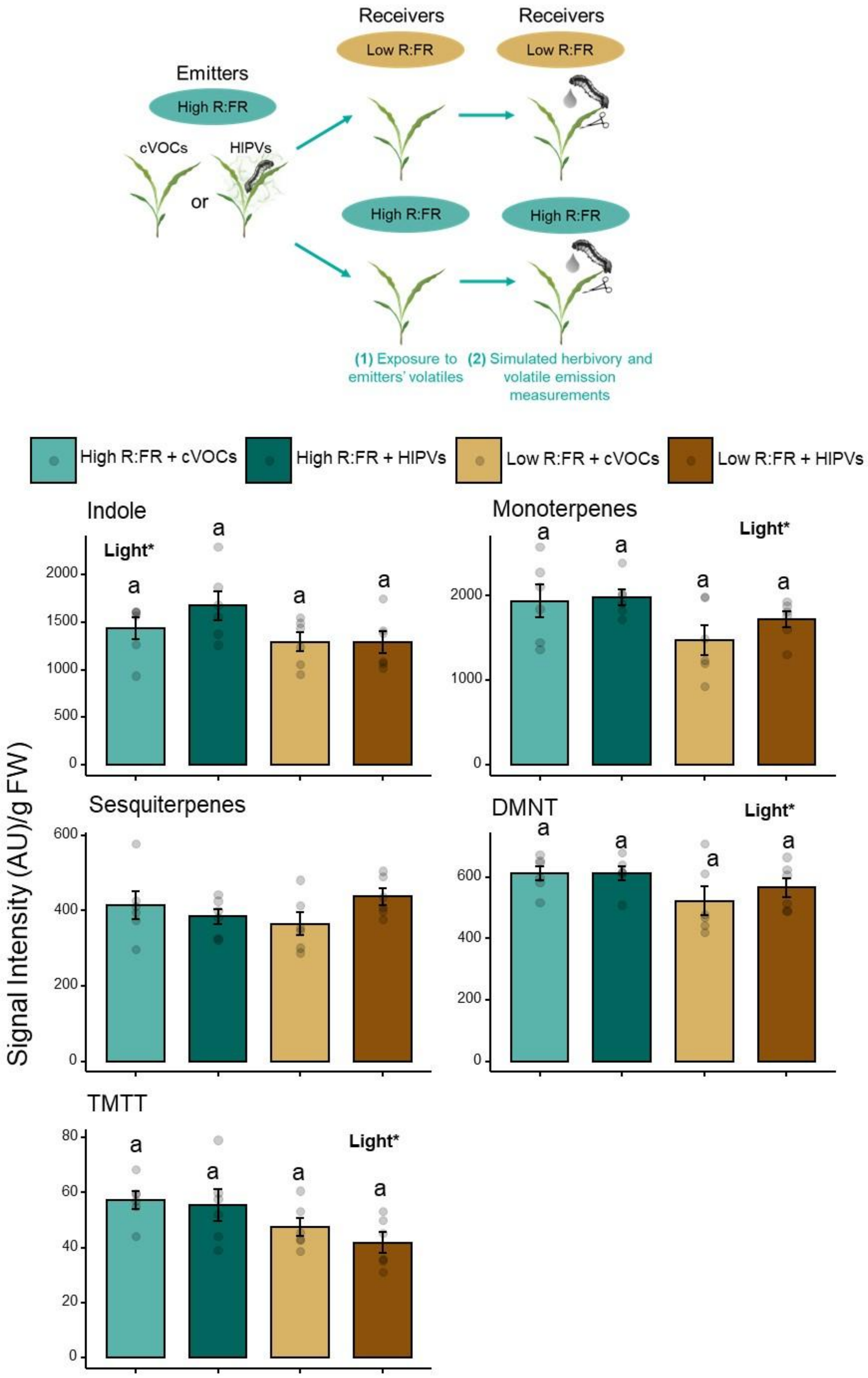


**Figure S6.** Sum (mean ± SEM, *n* = 6) of the emission of individual volatiles signatures measured over a period of 12 h upon simulated herbivory induction in receiver plants. Low or high R:FR-treated receiver plants were pre-exposed to cVOCs or HIPVs from high R:FR-treated plants during 20 h followed by simulated herbivory (wounding + oral secretions from *S. littoralis*).The effects of light treatment of the receiver (Light), the volatiles from the emitters (Vol), and their interactions on receivers' volatile emissions were tested using linear models followed by the calculation of estimated marginal means (EMMs) and pairwise comparisons using Tukey's Honest Significant Difference (HSD) test. Statistically significant effects are shown in the graph (*$p < 0.05$). Different letters denote significant differences among groups at $p < 0.05$. DMNT and TMTT stand for the homoterpenes (*E*)-4,8-dimethyl-1,3,7-nonatriene and (*E*, *E*)-4,8,12-trimethyltrideca-1,3,7,11-tetraene, respectively. AU refers to arbitrary units.
